# A novel function of *M. tuberculosis* chaperonin paralog GroEL1 in copper homeostasis

**DOI:** 10.1101/766162

**Authors:** Mohammed Yousuf Ansari, Sakshi D. Batra, Hina Ojha, Ashish, Jaya S. Tyagi, Shekhar C. Mande

## Abstract

Mycobacterial GroELs namely GroEL1 and GroEL2 belong to the family of molecular chaperones, chaperonins. Chaperonins in *Escherichia coli* are termed as GroEL and GroES which are encoded by essential genes and are involved in cellular protein folding. GroEL1 has a characteristic Histidine-rich C-terminus contrary to its essential paralog GroEL2 and *E. coli* GroEL which have hydrophobic (GGM) repeats. Since Histidine richness is likely to be involved in metal binding, in this study we have attempted to decipher the role of GroEL1 protein in chelating metals and the consequent role on *M. tuberculosis* physiology. Using isothermal titration calorimetry (ITC), we found that GroEL1 binds copper, nickel and cobalt, with the highest binding affinity to copper. Since copper is known to be toxic at higher concentration, we cultured Wild Type *M. tuberculosis* H37Rv, *groEL1* knock-out and *groEL1*-complemented strain with increasing concentrations of copper. We found that *M. tuberculosis groEL1* knock out strain is more sensitive to copper than the wild type. Further hypothesizing that the probable mode of action of copper is by induction of oxidative stress, we attempted to understand the role of GroEL1 in redox silencing and hydroxyl radical mediated DNA damage. We interestingly found through our *in vitro* studies that GroEL1 is helpful in protection from copper stress through maintaining redox balance and free radical mediated DNA damage. Thus, these results indicate that the duplication of chaperonin genes in *M. tuberculosis* might have led to their evolutionary divergence and resulted in a functional divergence of chaperonins.

## Introduction

Tuberculosis is a deadly disease caused by *Mycobacterium tuberculosis*, an acid-fast bacterium. Nearly 1.6 million people die of *M. tuberculosis* infections worldwide (WHO report, 2018). *M. tuberculosis* is transmitted from an infected person to a healthy person through aerosols by coughing or sneezing. The primary sites of infection are lungs, where the bacteria are phagocytosed by alveolar macrophages.

Most of the bacteria are killed by phagocytic machinery of macrophages (i.e avirulent *M. tuberculosis*), however virulent *M. tuberculosis* strains modulate the host defence mechanisms and survive inside the phagosome (1). Inside the phagosomes, mycobacteria encounter various stresses, such as low oxygen level, nutrient deficiency, reactive oxygen species and reactive nitrogen species (2). Mycobacteria have evolved elegant mechanisms to overcome these host responses for their survival in the host. Thus, in the interplay of host-pathogen interactions, both the host and the pathogen utilize multiple mechanisms and counter mechanisms to their advantage.

The concentration of inorganic ions are also known to be modulated inside the macrophages during an infection (3). For example, using hard X-ray microprobe, it was reported that the level of copper fluctuates between 25 µM to 500 µM inside the phagosomes infected with virulent *M. tuberculosis* (4). The levels of copper concentration have been shown to be increased within the granulomatous lesion in the lungs and lead to clearance of mycobacterial load in the guinea pig (5). In order to eliminate pathogen host modulates the concentration of copper ions inside infected phagosomes, however the mechanism is still not clear. The molecular basis by which the pathogen counters the Cu^2+^-ion (cupric ion) mediated defence of the host, is among the least understood processes.

The beneficial effects of copper on humankind have been known since antiquity and its antimicrobial property as well as other physiological functions are beginning to be understood in the recent era (6, 7). It has been reported that water stored in a copper pot is effective in killing major pathogens such as *E. coli, Shigella flexneri, Vibrio cholera, Salmonella typhi* and Rotavirus based on the ancient Indian medical practice, *Ayurveda* (8, 9). Early studies have shown that the efficacy of the drug isoniazid is enhanced when used in combination with copper (10, 11). The mode of action of copper on the physiology of *M. tuberculosis* is attributed to its high redox potential (12). It can exist in two oxidation states, Cuprous (Cu^1+^) and Cupric (Cu^2+^). Under aerobic conditions, copper has the tendency to undergo Fenton and Haber-Weiss reactions to generate hydroxyl radicals (13). Such hydroxyl radicals can cause damage to lipids, nucleic acids and proteins. The damage to proteins can be caused by an attack on the polypeptide backbone and disrupting their covalent structure or by attacking amino acids, such as cysteine, which can displace iron from the iron-sulphur cluster containing proteins (14, 15). Copper may be involved in these reactions causing further damage (16). To resist the effect of copper toxicity, *M. tuberculosis* might employ various mechanisms such as copper efflux by P-type ATPases; utilizing oxidases to convert toxic cuprous ion to less toxic cupric ion; or sequestrating copper to maintain the level of copper ions (17–19).

We recently reported that *M. tuberculosis groEL1*-KO (knockout) strain is significantly sensitive to growth under low-aeration stress condition (20). Moreover, it was intriguing to note that nine of the thirty copper responsive genes (20), are downregulated during the growth of the *groEL1*-KO strain. Copper responsive genes were identified by culturing *M. tuberculosis* under physiological levels of copper in the culture media. It was shown that high level of copper is toxic for the survival of *M. tuberculosis* and it was also reported that oxidative stress response genes were induced under high concentration of copper in culture media (21). Oxidative stress response is also the manifestation of low aeration stress to *M. tuberculosis* (20). We therefore wished to test the hypothesis that GroEL1 of *M. tuberculosis* might be involved in resistance to Cu-toxicity. In this present study we have attempted to dissect the role of GroEL1 protein at the interface of *M. tuberculosis* and copper stress.

*M. tuberculosis* possesses two copies of *groEL* genes, *groEL1* and *groEL2*. *groEL1* is arranged in an operon along with cognate co-chaperonin *groES* while the *groEL2* gene exists separately on the genome (21, 22). Moreover, unlike groEL1, *groEL2* is essential for survival for *M. tuberculosis* in host. A phenotype of granuloma formation defect in guinea pig when infected with *M. tuberculosis groEL1* knockout strain has been reported (23). Evolutionary studies on *M. tuberculosis groEL* sequences have shown rapid evolution of the gene and adapting novel properties different from the basic protein folding function (24). One of the distinct properties is their failure to form canonical tetra-decameric structure and exist as lower oligomers when purified recombinantly from *E.coli* (25). However, when purified from *M. tuberculosis* culture filtrates, GroEL1 was found to exist in different oligomeric forms such as dimer, heptamer and tetradecamer. It was also shown that the tetra-decameric form is phosphorylated (26). Interestingly, *M. tuberculosis* GroEL1 is devoid of Gly-Met rich C-terminal tail which is known to be crucial for chaperoning function (27) but instead possesses a Histidine-rich C-terminal region. Since the Histidine-rich sequences are known to bind cations, we have attempted to explore the role of GroEL1 in copper-binding if any, and the consequent physiological effects.

We show that GroEL1 indeed binds copper ions and that the *groEL1*-KO strain of *M. tuberculosis* is susceptible for higher copper ion concentrations as compared to the Wild Type strain. We also show that the killing mechanism could be due to the imbalance of redox state and hydroxyl radical mediated DNA damage. The present study on GroEL1 protein has revealed a novel functional property of copper sequestration, a function that was unknown in *M. tuberculosis*.

## Results

### Sequence analysis and homology model of GroEL1

To understand the structural aspects of GroEL1, we modelled GroEL1 protein on the basis of crystal structure of its paralog GroEL2 and SEC-MALS data indicating that it is a monomer in solution. Homology model of *M. tuberculosis* GroEL1 showed expectedly that it can be divided into three domains – Apical (189-374 residues), Intermediate (132-188 and 375-406 residues) and Equatorial domain (1-131 and 407-539 residues). Multiple sequence alignment of *E. coli* GroEL, *M. tuberculosis* GroEL1 and GroEL2 reveal that they have distinctive C-terminal amino acid sequences. Contrary to *E. coli* GroEL and *M. tuberculosis* GroEL2, which possess (GGM) repeats, *M. tuberculosis* GroEL1 has a Histidine-rich C-terminal (Figure 1A). In the homology model, N and C-terminal regions, due to low complexity sequences, were expectedly found unstructured (Figure 1B). In the tetradecameric structure of chaperonins, these residues protrude in the central cavity of the tetradecamer and are known to play a role in protein folding function of the chaperonins.

**Figure 1:**
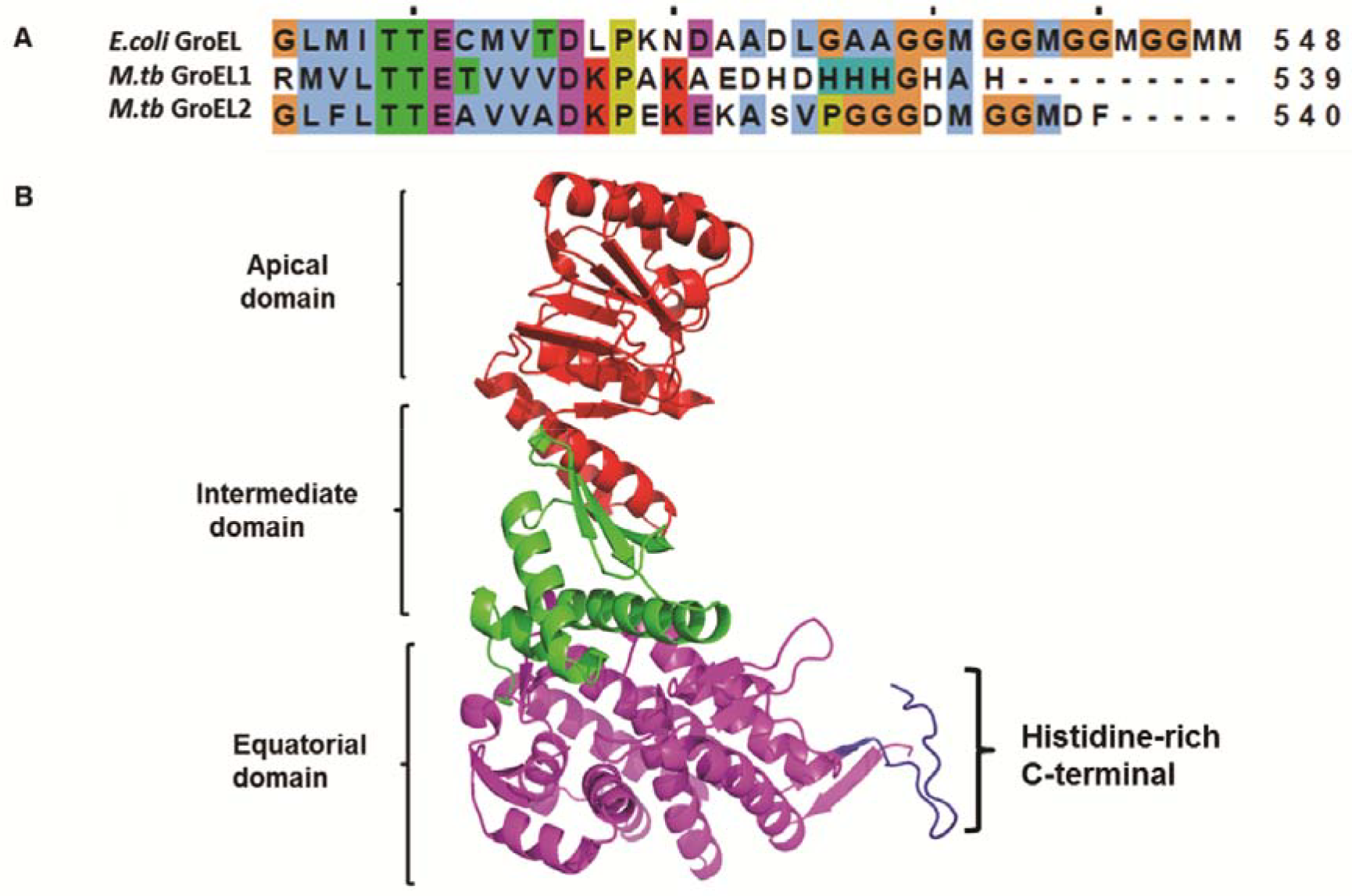
Sequence analysis and homology model of GroEL1. **(A)** Gene sequences of *E. coli* GroEL, *M. tuberculosis* GroEL1 and GroEL2 were analysed (Jalview) using multiple sequence alignment program Clustal Omega (EMBL-EBI). Contrary to essential chaperonins i.e *E. coli* GroEL and *M. tuberculosis* GroEL2, having (GGM)4 repeats, *M. tuberculosis* GroEL1 has Histidine-rich C-terminal. **(B)** Homology model of *M. tuberculosis* GroEL1 was built using *M. tuberculosis* Cpn 60.2 crystal structure, 2.8 Å (PDB, 3RTK A). The software used was Modeller v9.18. It can be divided in three domains – apical, intermediate and equatorial domain (1-133 and 409-539 residues). N and C-terminal region were found unstructured in the model. The last 18 residues from C-terminal (purple) were deleted for biophysical studies.

### GroEL1, GroEL1-Δ18 and GroEL1-ED exists as monomer

For understanding the functional role of *M. tuberculosis* GroEL1, its 18 C-terminal residues deleted mutant (GroEL1-Δ18), and the equatorial domain (GroEL1-ED) were recombinantly expressed and purified from *E. coli*. Size exclusion chromatography coupled with multi-angle light scattering (SEC-MALS) of all the three purified proteins showed that the proteins eluted as single peaks. However, the light scattering profile showed two peaks, the first peak showed higher soluble aggregates whereas the second peak was consistent with the theoretical molecular weight of a monomer *i.e*. ~56 kDa (Figure 2). The results were consistent with the full length GroEL1 and GroEL1-∆18 proteins being monomers (Figure 2). The purified GroEL1-ED showed that the molecular weight is closer to its estimated theoretical molecular weight of a monomer i.e. ~27 kDa (Figure S1).

**Figure 2:**
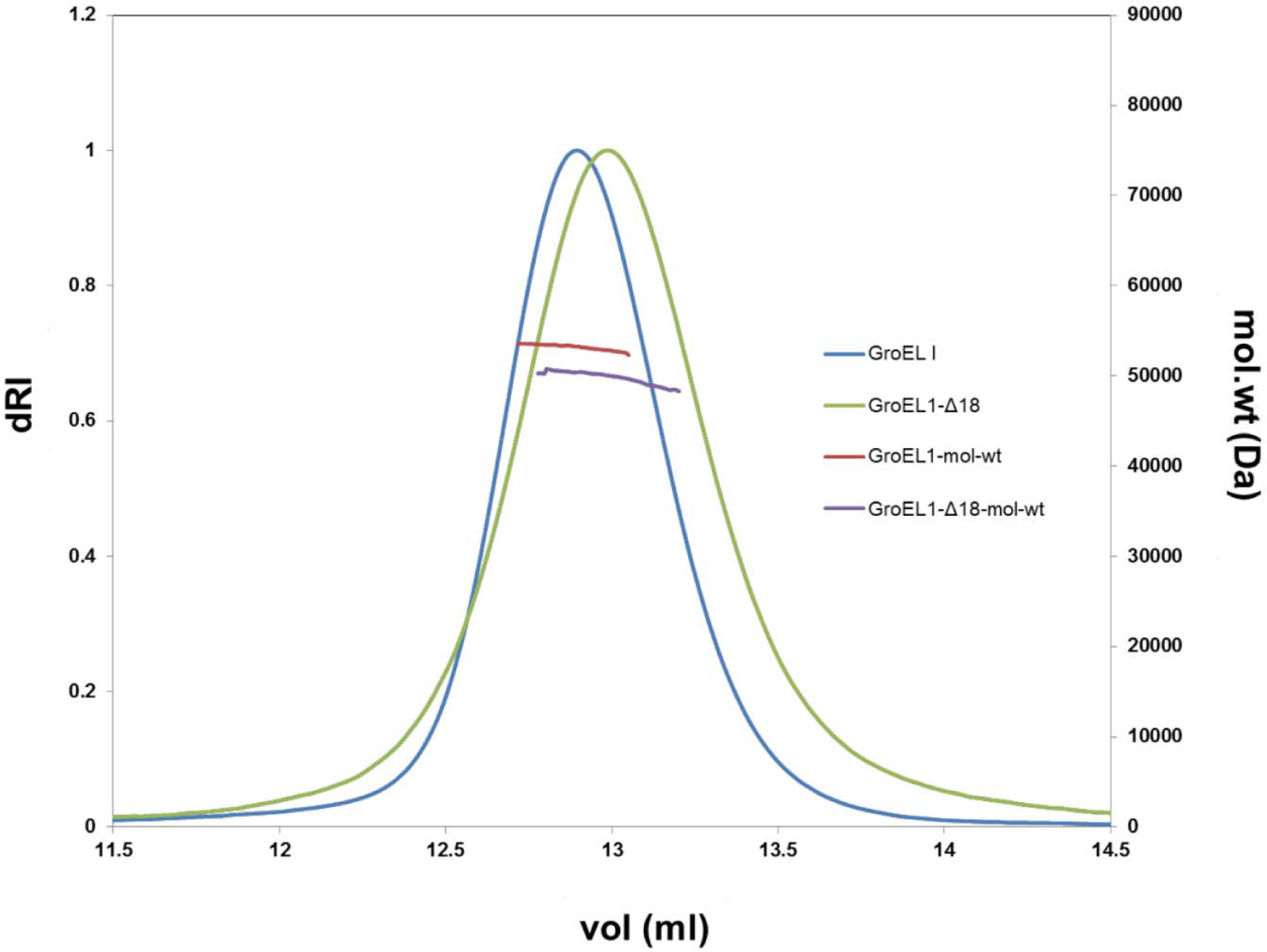
GroEL1 and GroEL1-Δ18 exists as monomer as shown by SEC-MALS. SEC-MALS elution profiles of GroEL1 and GroEL1-Δ18 in 10 mM Tris-Cl pH 8.0 and 150 mM NaCl at 1.5 mg/ml concentration respectively. The molecular mass distributions are shown as thick continuous lines. The data was analysed using Astra software.

### GroEL1 binds Copper (Cu^2+^) and other divalent transition metals

Data analysis from isothermal titration calorimetry (ITC) showed that copper (Cu^2+^) (Figure 3A), nickel (Ni^2+^) (Figure 3B) and cobalt (Co^2+^) (Figure 3C) bind to GroEL1. We fitted nickel and cobalt to one binding site model where both the binding isotherms were observed to be exothermic. The reactions had Gibbs free energy ΔG, ~-27 kcal/mol and ~-30 kcal/mol respectively, and the binding affinity was found to be in micro-molar range (ΔKd ~17 µM and ~6 µM, respectively). The binding isotherm of copper showed exothermic (molar ratio is ~1.5) followed by endothermic pattern (molar ratio is ~3). We fitted the isotherm in ‘sequential two binding model’ with the two binding sites of Gibbs free energy ΔG, ~-33 kcal/mol and ~-19 kcal/mol, respectively). The first site has a better binding affinity (ΔK_d_ is ~2 µM), whereas the second site has a lesser binding affinity (ΔK_d_ is ~441 µM). Titration with other transition metals did not show characteristic binding isotherm. We also tried to probe interaction with iron as it has been hypothesized to interact with GroEL1 (28). Titration with ferric ion did not show interaction (Figure 3D). Thus, the ITC results indicate that GroEL1 binds copper (Cu^2+^) with higher affinity compared to nickel and cobalt (Table 1).

**Table 1:** Thermodynamic parameters for metal binding to GroEL1 determined by ITC. The table shows thermodynamic parameters obtained after respective model fitting for nickel, cobalt and copper ions binding to GroEL1 using MicroCal ITC200 Origin software. The experiments were performed at 25°C. Standard deviation shown is from three biological replicates. N is the binding stoichiometry.

|  | Nickel | Cobalt | Copper |
| --- | --- | --- | --- |
| **Binding model** | **ONE** | **ONE** | **SEQUENTIAL TWO** |
| N | 1.5±0.42 | 2.97±0.95 |  |
| $\Delta K1 (M^{-1})$ | 6.03E+04±1.42E+04 | 1.82E+05±8.30E+04 | 9.61E+05±8.57E+05 |
| $\Delta H1 (cal/mol)$ | -1.66E+04±1.95E+03 | -6.95E+03±2.50E+03 | -3.11E+03±1.25E+03 |
| $\Delta S1 (cal/mol/deg)$ | -3.39E+01±6.10 | 0.64±8.55E+00 | 1.64E+01±5.90E+00 |
| $\Delta K2 (M^{-1})$ | | | 2.51E+03±8.94E+02 |
| $\Delta H2 (cal/mol)$ | | | 4.10E+04±3.21E+04 |
| $\Delta S2 (cal/mol/deg)$ | | | 152.97±106.77 |
| $\Delta Kd1 (M)$ | **1.71E-05±3.65E-06** | **6.19E-06±2.35E-06** | **1.69E-06±1.19E-06** |
| $\Delta Kd2 (M)$ | | | **4.41E-04±1.81E-04** |
| $\Delta G1 (cal/mol)$ | -2.72E+04±5.60E+02 | -2.99E+04±1.06E+03 | -3.35E+04±2.18E+03 |
| $\Delta G2 (cal/mol)$ | | | -1.93E+04±9.71E+02 |

**Figure 3:**
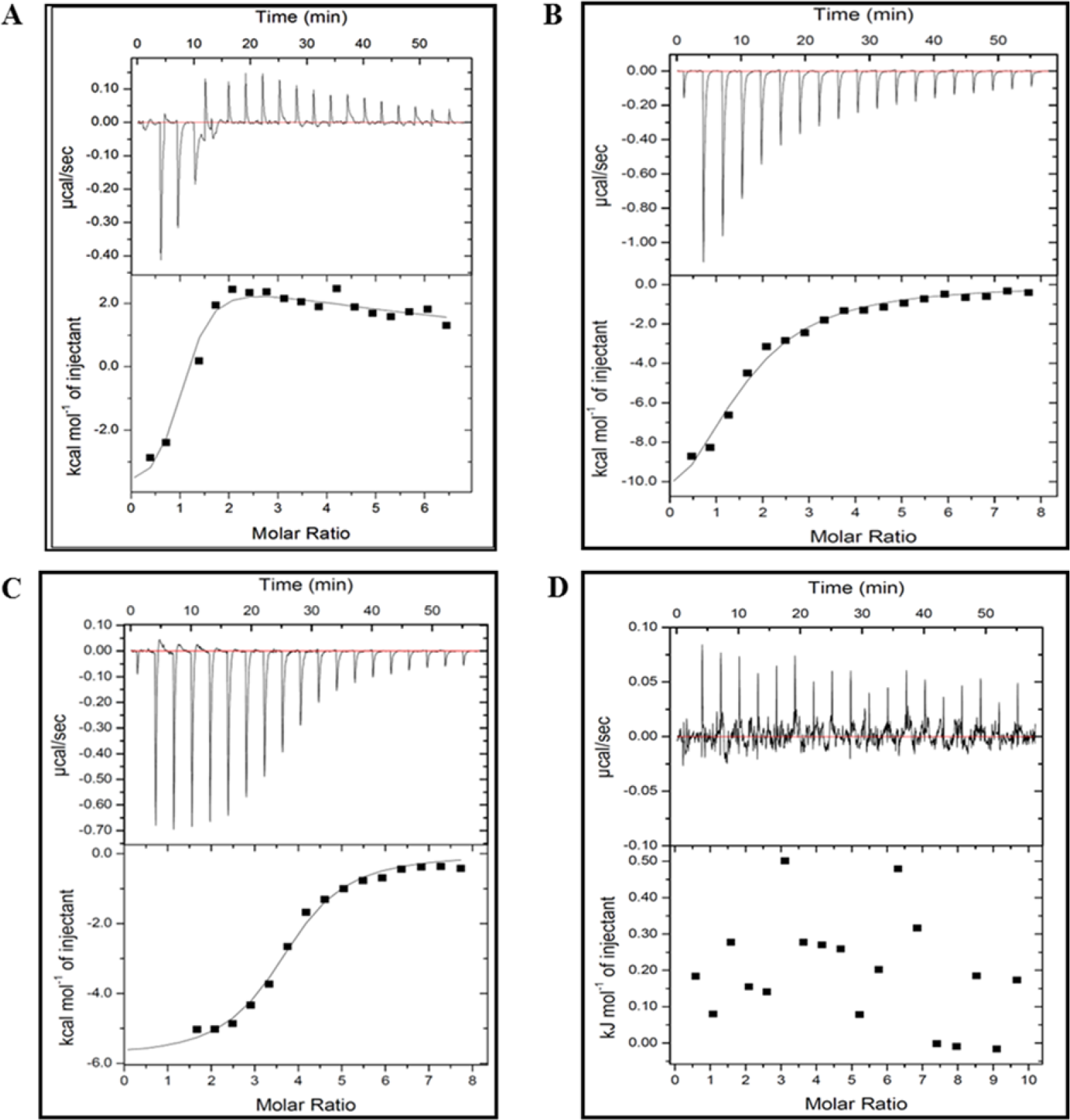
GroEL1 binds copper as shown by ITC. Chloride salts of **(A)** Cupric chloride (1mM) **(B)** Nickel chloride (1mM), **(C)** Cobalt chloride (1mM) and **(D)** Ferric chloride (1mM) were titrated into GroEL1 (~30µM) in 10 mM Tris-Cl pH 8.0. In the representative figure the upper panel shows heat peaks after consecutive titration of salt ligands into GroEL1 after subtracting heat of dilution generated by titrating in the buffer alone. The lower panel shows integrated heat data fit to sequential two binding site model for **(A)**, one site model for **(B)** and **(C)**. Data points for ferric ions (Fe^2+^) show no binding. **Refer** Table 1 for thermodynamic parameters obtained after model fitting.

### Histidine-rich C-terminus of GroEL1 protein is essential binding site for Copper (Cu^2+^)

To confirm the involvement of Histidine-rich C-terminal in metal binding, we deleted the last 18 residues (GroEL1-Δ18) and performed ITC. As expected, binding to Cu^2+^, Ni^2+^ and Co^2+^ was lost in the C-terminal deleted protein (Figure 4A). As the data points were noisy we could not derive any thermodynamic parameters. These results clearly indicate that the last 18 residues are essential for metal binding.

**Figure 4:**
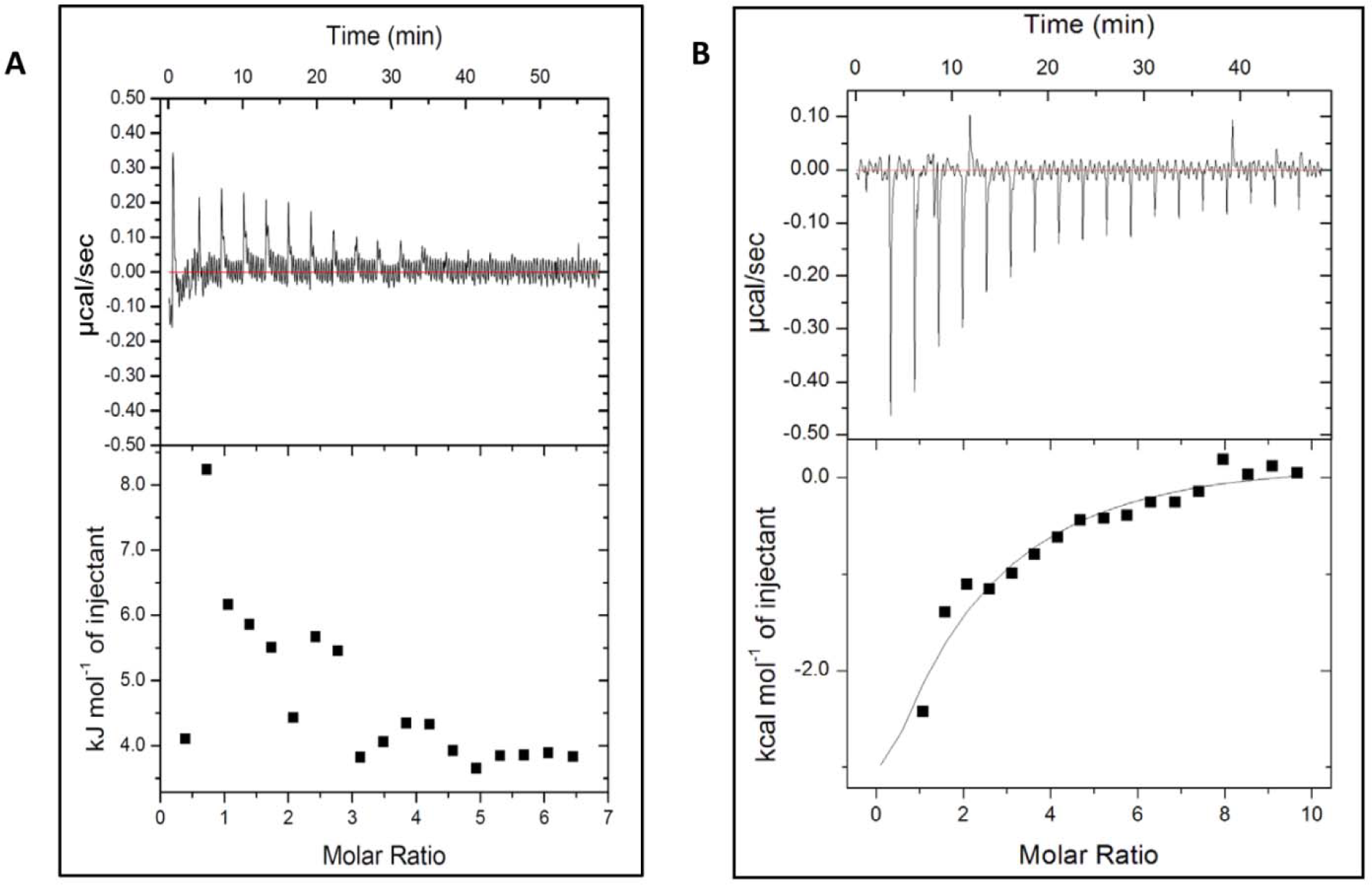
Histidine-rich C-terminus of GroEL1 protein is essential binding site for Copper.. **(A)** The chloride salt of cupric chloride (1mM) was titrated into the ITC cell containing ~30µM GroEL1-Δ18 in 10 mM Tris-Cl pH 8.0. In the representative figure, the heat generated through successive titration indicates endothermic binding curve but the data fitting is very noisy. Data fitting is not suitable for estimating thermodynamic parameters. **(B)** Chloride salt of cupric chloride (1mM) was titrated into GroEL1-ED (30µM) in 10 mM tris-Cl pH 8.0. In the representative figure the upper panel shows heat peaks after consecutive titration of copper into GroEL1-ED after subtracting heat of dilution generated by titrating copper in the buffer alone. The lower panel shows integrated heat data fit to one binding site model.

As the C-terminal residues are a part of the equatorial domain of GroEL1, we performed ITC with the purified equatorial domain of GroEL1. Similar metal binding was observed for copper (Cu^2+^) ions by the equatorial domain as the full length protein. However, the endothermic isotherm seen in copper (Cu^2+^) ions binding to full length GroEL1 was lost. There was only exothermic binding and we were able to fit the isotherm to ‘single binding site model’ (Figure 4B). The result suggests that the second weak binding site for Cu^2+^ ion is not present or related to the equatorial domain.

### Copper (Cu^2+^) ions cause large scale conformational change in GroEL1

To understand the mechanistic flow, shape characterization of GroEL1 was explored by Small Angle X-ray Scattering (SAXS) experiments. The double log plot **(Log_10_ I(Q) versus Log_10_ Q)** of each dataset (GroEL1 and its complex with Cu^2+^ ions) confirmed no upward or downward change in the low Q-region of the intensity profile reminiscent of aggregation or inter-particle interference respectively (29) (Figure 5A, Figure S3). Linearity of the Guinier analysis considering globular nature of scattering particles further confirmed monodisperse profile of the sample during data collection (**Q^2^ versus Ln I(Q)**, Figure 5B, Figure S3).

**Figure 5:**
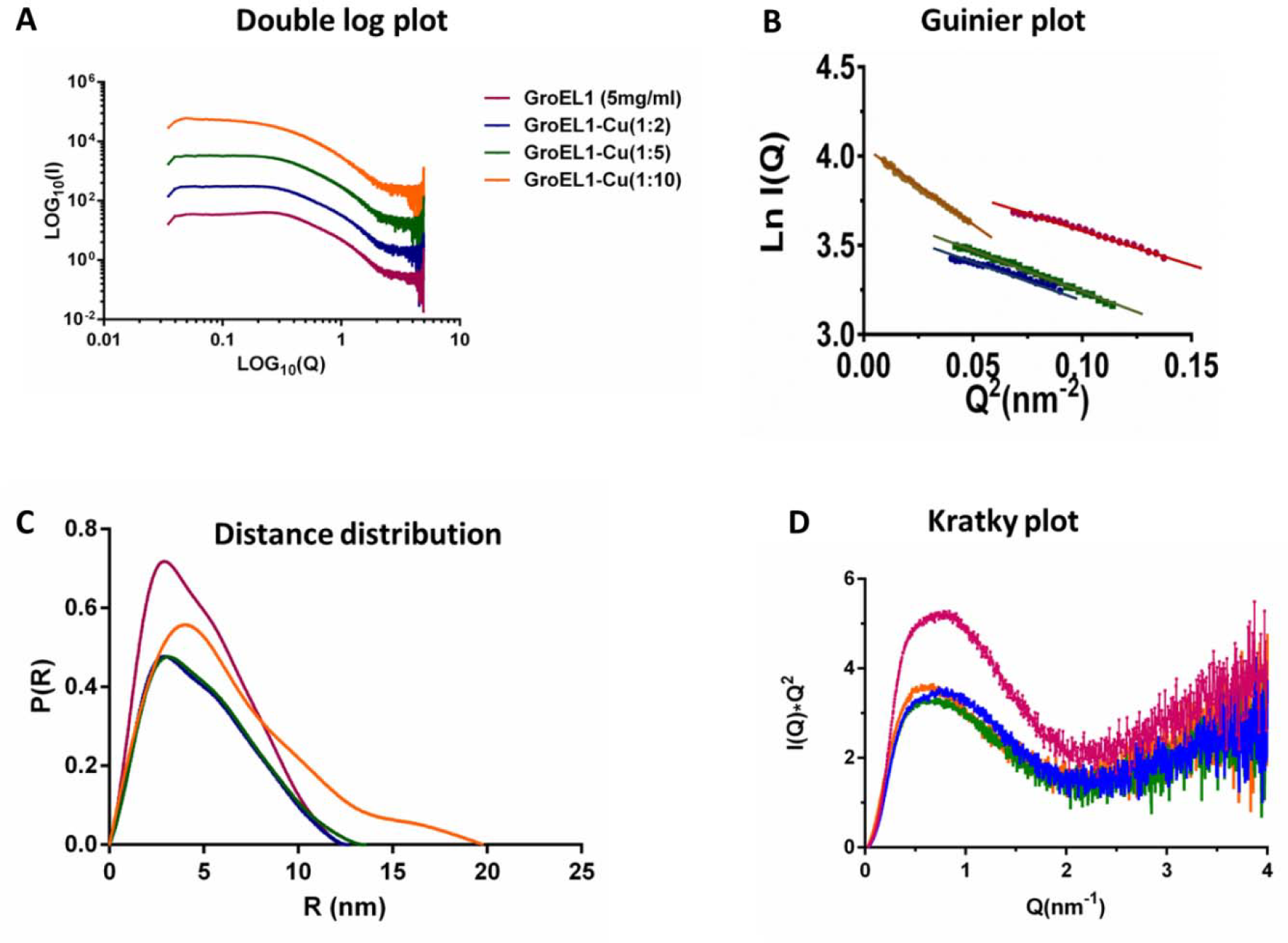
SAXS data analyses and plots of GroEL1 with copper (Cu^2+^) SAXS data analysis of GroEL1 protein in complex with Cu^2+^ ions (CuCl_2_) in the molar ratio 1:2, 1:5 and 1:10. (A) Double-Log plot of SAXS data of GroEL1 and Cu^2+^ ions (CuCl_2_) complex (data curve is modified for easy visualisation). (B) Guinier plot of GroEL1 and Cu^2+^ ions (CuCl_2_) complex (data curve is modified for ease in visualisation). (C) Pairwise distance distribution plot of GroEL1 and Cu^2+^ ions (CuCl_2_) complex. (D) Kratky plot of GroEL1 and Cu^2+^ ions (CuCl_2_) complex.

Concentration dependent characterization of GroEL1 protein suggests that there was no change in R_g_ value derived from Guinier analysis. GroEL1 at 5 mg/ml and 2.5 mg/ml showed similar R_g_ values of 3.32 nm and 3.21 nm respectively. The R_c_ remained constant at approximately 1.7 nm, suggesting no aggregation of proteins. D_max_ as was obtained from computing pairwise distance distribution curve P(r) from the two concentration range (i.e. 5 mg/ml and 2.5 mg/ml) using indirect Fourier transformation. It was estimated to be 12.49 nm and 12.36 nm respectively and was comparable to the estimated persistence length (L) of molecules using only low Q data. The hyperbolic profile of the Kratky plot of each dataset suggests Gaussian chain-like shape in the solution. There was no change in the peak position (Figure S3, Table 3).

Further SAXS studies of GroEL1 (5mg/ml) in presence of copper (Cu^2+^) at molar ratio of 1:2, 1:5 and 1:10 show that the R_g_ values derived from Guinier analysis increased from 3.32 to 5.07 nm of GroEL1 protein in the presence of Cu^2+^ ions, however the R_c_ values remained constant at approximately 1.7 nm (Table 3). The D_max_ values were found to be comparable to the estimated persistence length, L of molecules using only low Q data. There was an increase in the D_max_ values of GroEL1 protein in the presence of Cu^2+^ ions from 12.49 to 19.75 nm (Figure 5C, Table 3). The Kratky plot shows change in peak position in the presence of copper (Cu^2+^) ions correlating with the increase in particle size (Figure 5D, Table 3). The Porod-Exponent value (measure of extent of flexibility in scattering molecule) obtained from each dataset implies that GroEL1 structure opens-up in the presence of copper (Cu^2+^) ions and becoming more flexible (30). The Porod-Exponent value decreases gradually from 3.5 to 3.2 (Table 3). These results indicate GroEL1 protein becoming larger in presence of copper (Cu^2+^) ions and inherent disorder or overall flexibility increases in the protein molecules with the addition of or binding to Cu^2+^ ions. Additionally, molecular weight of scattering particle in the sample was estimated based on a SAXS invariant V_c_ (volume of correlation), which is concentration-independent and specific to the structural state of the scattering molecule in solution (31). The apparent molecular weight value ranged from 55.3 to 80.1 kDa of GroEL1 protein and its complex with copper (Cu^2+^)ions, suggesting that GroEL1 protein remains monomer and a slight increase in hydrodynamic properties occurs (Molecular weight calculated from the sequence is 55.8 kDa) (Table 3).

The SAXS data was used to calculate the three-dimensional shape of GroEL1 protein and its complex with copper (Cu^2+^) ions. Being aware of the inherent disorder in the molecules, and limitation of uniform density modeling and averaging in interpreting such shapes, chain-ensemble modeling was done and one model was selected over others as described in methods section. The final χ^2^ values are reasonably good for the SAXS data based models (Table 3). Dummy atom models were superimposed on the homology model of GroEL1 protein in an automated manner. The superimposition revealed a large degree of similarity in the shape profile, however, there is extra volume near the two extremities in the SAXS-based models. This may be due to domain motions (Figure S4). A large scale structural rearrangement in GroEL1 with the opening-up of the structure in presence of copper (Cu^2+^) ions was also observed (Figure 6).

**Figure 6:**
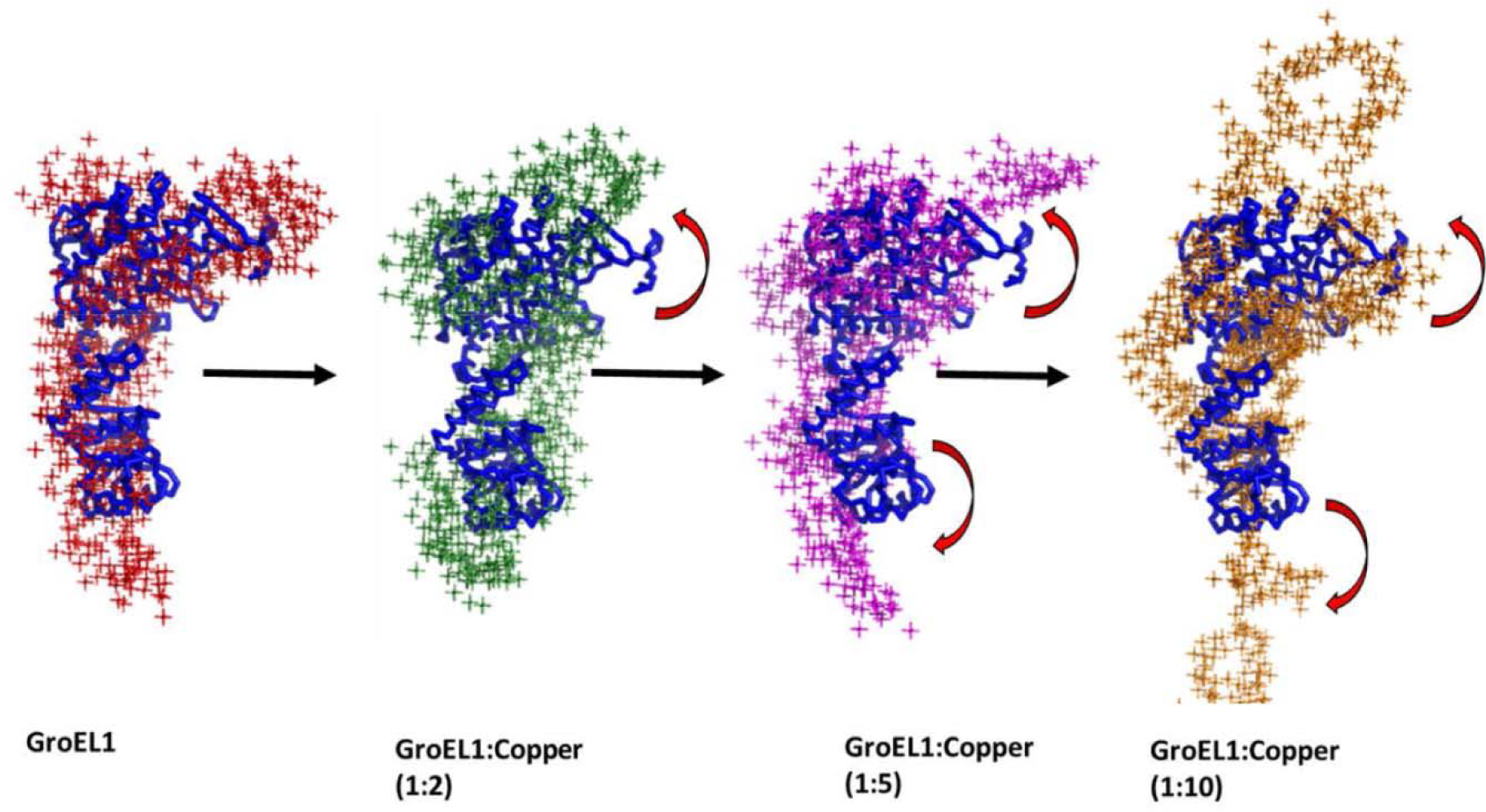
SAXS based *ab initio* models of GroEL1 with copper (Cu^2+^) Predominant solution shape of SAXS based model of GroEL1 and Cu^2+^ ions (CuCL_2_) complex in the molar ratio 1:2, 1:5 and 1:10. Conformational changes observed in *ab initio* model (GASBOR) of GroEL1 protein and its complex with Cu^2+^ ions (CuCL_2_). The models are superimposed on homology model of GroEL1 protein.

### Monitoring Copper (Cu^2+^) mediated conformation change in GroEL1 protein

In order to understand conformational change (shown through SAXS studies) mediated status of hydrophobic residues of GroEL1 protein, ANS (8-anilino-1-naphthalenesulfonate) assay was performed. ANS (8-anilino-1-naphthalenesulfonate) is a fluorescent dye that binds to the solvent-accessible hydrophobic residues in proteins. Conformation changes affecting solvent accessibility of hydrophobic residues can therefore be monitored by changes in ANS fluorescence intensity. The conformational change as observed in the SAXS experiments was also seen in GroEL1 (10 μΜ) in presence of copper (Cu^2+^) in the molar ratio of 1:5, 1:10 and 1:20 evident by the decrease in the intensity of ANS dye (Figure S2). A decrease in intensity could be manifested by the decrease in solvent-accessibility of hydrophobic residues of GroEL1 protein in the presence of copper (Cu^2+^) ions.

### Increased sensitivity of *M. tuberculosis groEL1*-KO (knockout) to copper (Cu^2+^)

In view of the observation that GroEL1 has a high affinity for copper (Cu^2+^) ions (Table 1), it was speculated that GroEL1 might have protective role for *M. tuberculosis* in the presence of high concentrations of copper (Cu^2+^) ions. Thus, the growth of *M. tuberculosis* strains *groEL1*-WT (wild type), *groEL1*-KO (knockout) and *groEL1*-comp (complemented)) was monitored in the presence of different concentrations of copper (Cu^2+^) ions (0, 5, 50 and 100 μM CuSO_4_) over a period of 16 days. There was nearly 2 to 2.5-fold decrease in the A_595_ of *groEL1*-KO (as compared to WT strain) in the presence of 100 μM CuSO_4_ from day 8 onwards (Figure 7), which was also in concordance with the viability assay (Figure 7). CFU (colony forming unit) results showed ~5.8 fold drop in the number of colonies obtained with *groEL1*-KO (as compared to WT strain) in the presence of 100 μM CuSO_4_ at day 16, indicating increased sensitivity of knock-out strain to high amount of copper (Cu^2+^) ions (Figure 7). We observed somewhat similar effect with 50 μM CuSO_4_ concentration (data not shown), however, 5 μM CuSO_4_ concentration was found to be little effective in CFU assay (data not shown). The significant drop in the *groEL1*-KO strain viability in the presence of copper (Cu^2+^), highlighted the plausible protective role of GroEL1 against copper toxicity.

**Figure 7:**
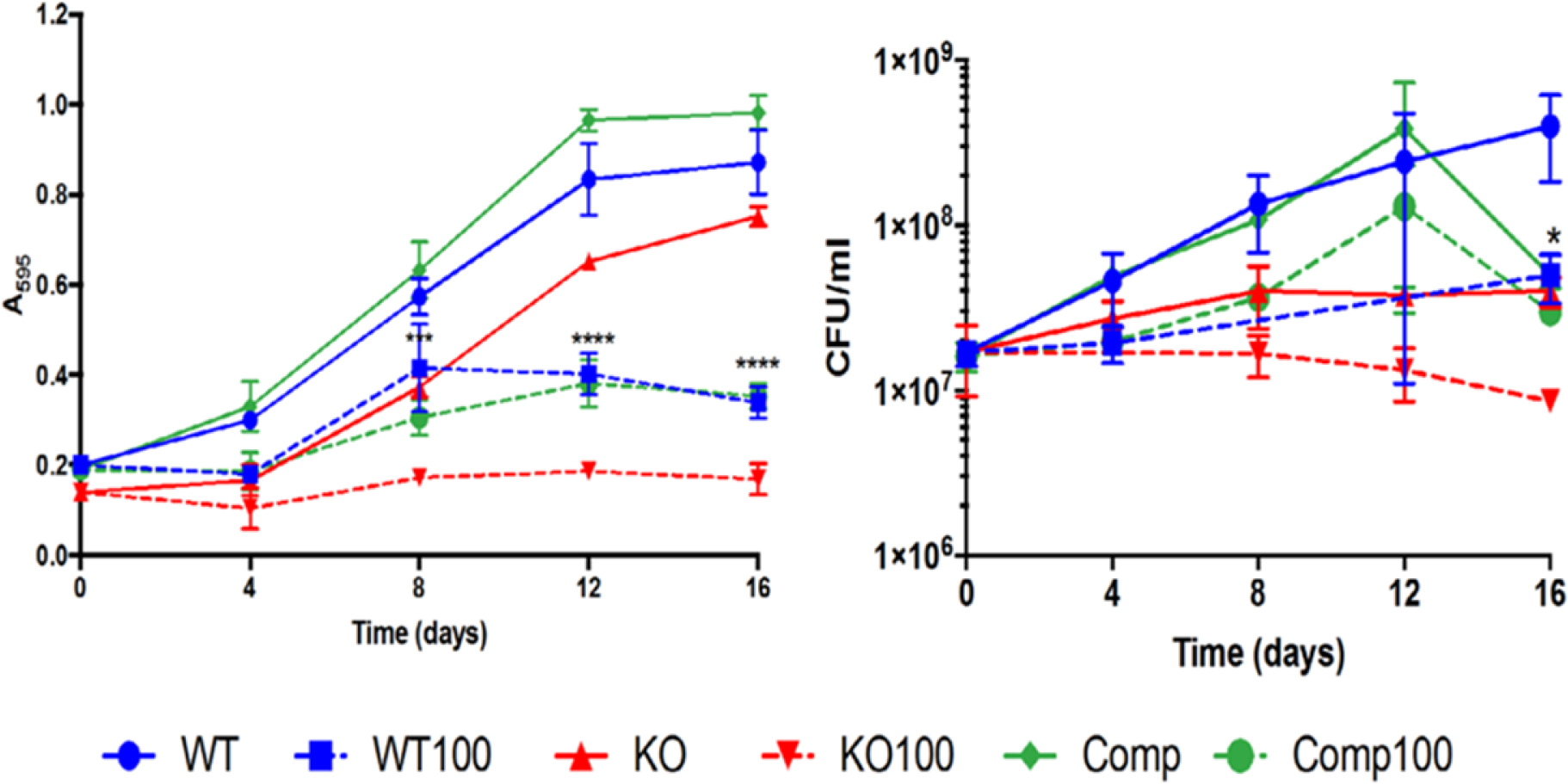
Copper tolerance in the *M. tuberculosis* GroEL1-KO strain is decreased. Figure shows the A_595_ and CFU/ml of WT (wild type), KO (knockout) and Complemented strains of *M. tuberculosis* at CuSO_4_ concentration 0 and 100 μM. Bold lines represent untreated (0 CuSO_4_) and dotted lines represent 100 μM CuSO_4._ Error bars are calculated from n= 3 to 6. ****<0.0001, *<0.05 calculated between WT100 and KO100 at respective time-points using unpaired t-test.

### Redox silencing by GroEL1 *in vitro*

Copper, like other transition metals, is involved in redox physiology in different organisms (32). Since GroEL1 binds copper (Cu^2+^) ions, it might modulate redox balance (33). We employed ascorbic acid as a model substrate to study redox silencing by GroEL1. Oxidation of ascorbic acid in the presence or absence of CuCl_2_ was observed spectrophotometrically at 260 nm during the span of 30 minutes. Ascorbic acid oxidized completely in the presence of CuCl_2_ while in absence of CuCl_2_ there was not much change in oxidation. When we added GroEL1 in the same reaction mixture containing Ascorbic acid and CuCl_2_, oxidation was decreased (Figure 8). GroEL1-Δ18, which was shown not to bind cations, was used as the control. The result indicates that GroEL1 is involved in preventing oxidation of ascorbic acid and hence, modulating redox balance *in vitro*.

**Figure 8:**
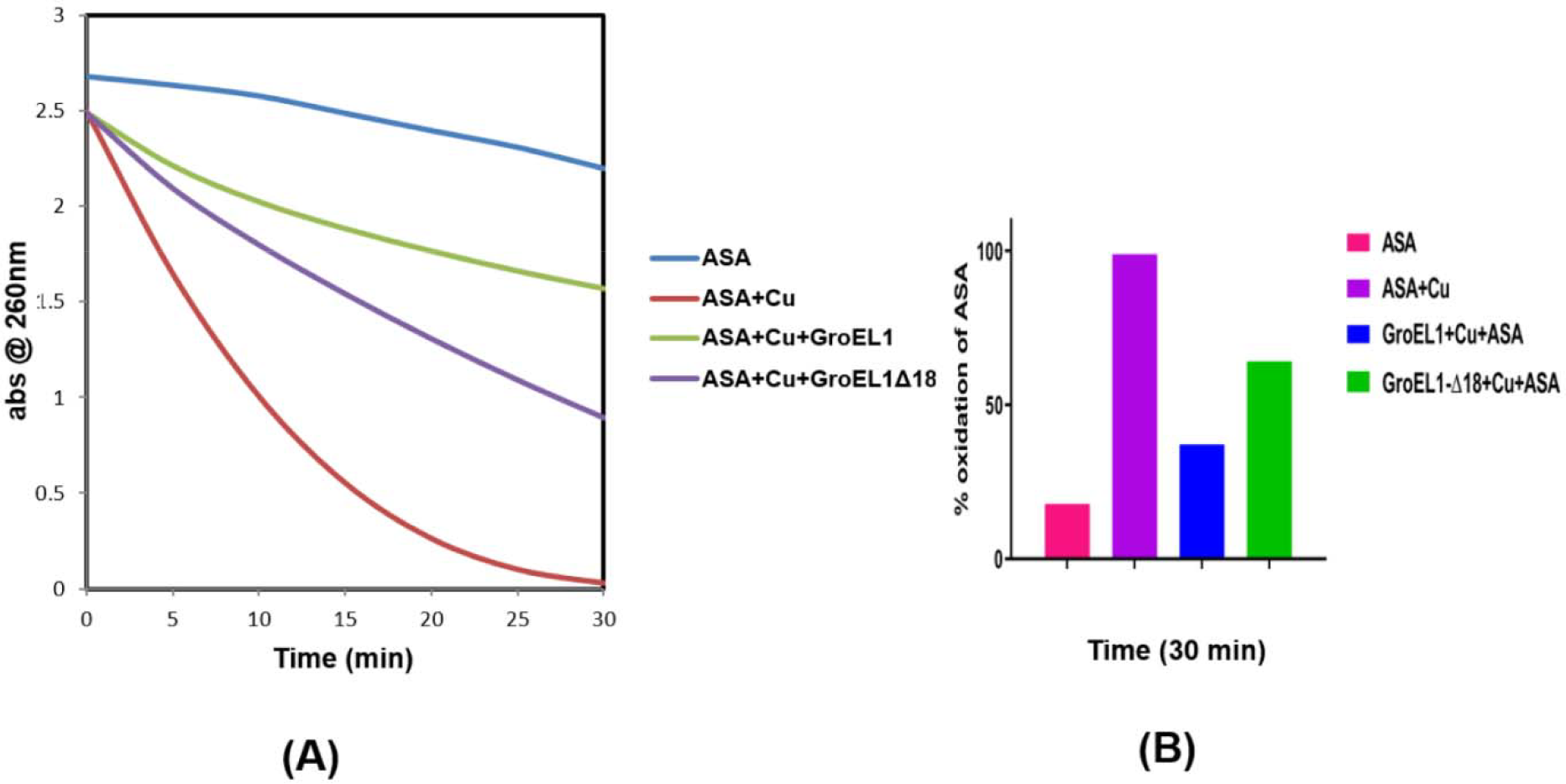
Cu^2+^-mediated ascorbic acid (ASA) oxidation system. The assays were performed using 300µM ASA, 5µM CuCl_2_ and absence or presence of 5µM GroEL1/GroEL1Δ18 at room temperature. The absorbance of ascorbic acid was measured at 260 nm during the span of 30 minutes. Oxidation of ascorbic acid in the presence cupric ion reduces the absorbance with time. Effect of GroEL1 on ^2+^-induced oxidation of ascorbic acid (ASA), suggest that it is involved in oxidation dependent redox balance (A). Bar graph showing percent oxidation of ASA from best of three replicates (B).

### DNA damage protection by GroEL1 *in vitro*

Copper follows Fenton-type of reaction to generate hydroxyl radicals in the presence of ascorbic acid and hydrogen peroxide (34). These hydroxyl radicals are effective in DNA damage. Since GroEL1 known to bind mycobacterial nucleoid (35), we tried to analyze its role in protecting copper catalyzed DNA damage. In the reaction mixture containing pBSK plasmid DNA, cupric chloride (CuCl_2_), ascorbic acid and hydrogen peroxide, there was complete plasmid DNA damage after incubating for 10 min at room temperature. This was observed on agarose gel (Figure 9). This phenomenon was lost when we added GroEL1 in the reaction mixture before adding Ascorbic acid and Hydrogen peroxide in the reaction mixture. We used GroEL1-Δ18 and *E. coli* GroEL as controls, where it fails to protect DNA damage. The result indicates that GroEL1 is instrumental in protecting DNA from copper catalyzed hydroxyl radical formation.

**Figure 9:**
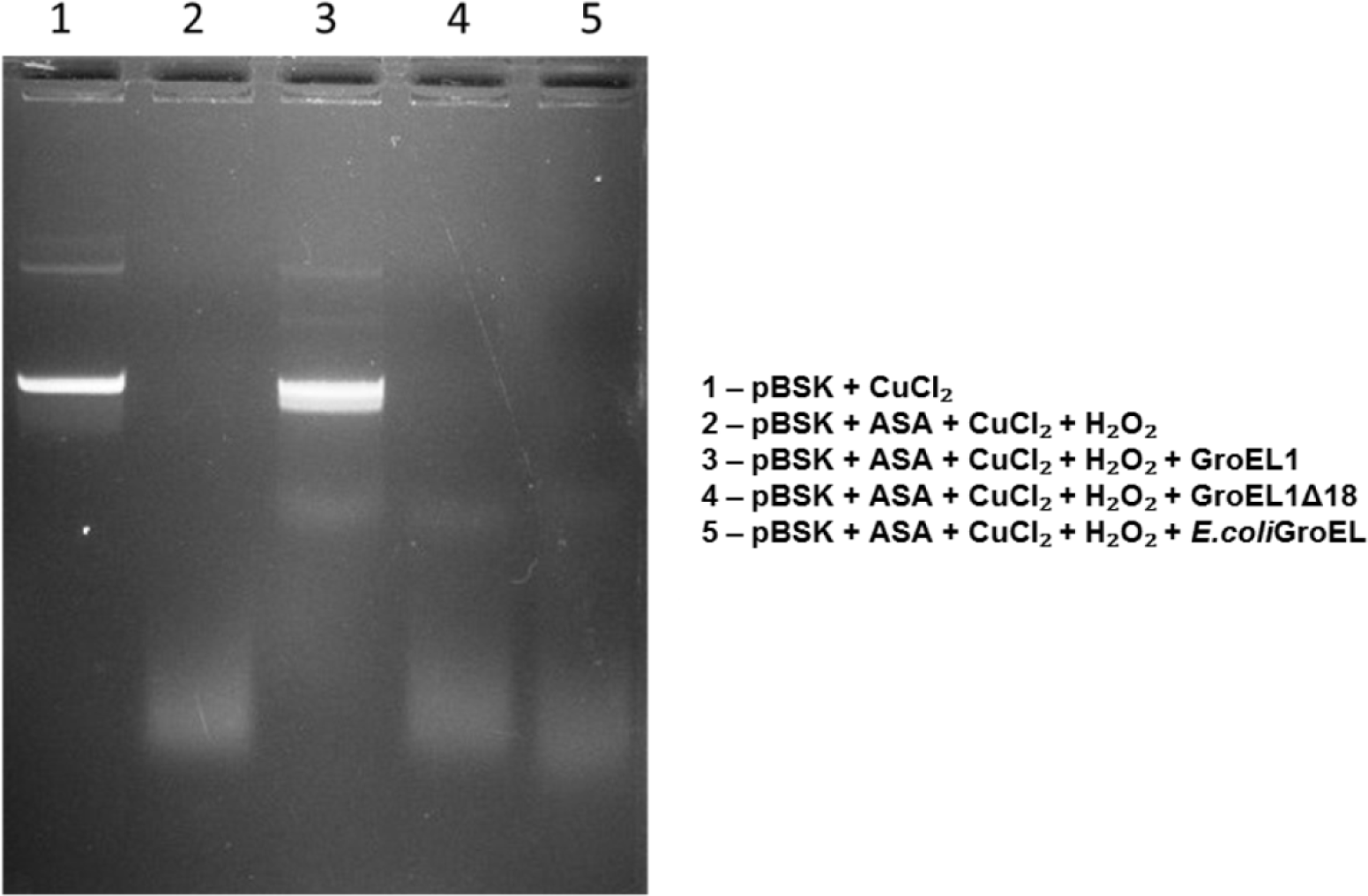
Protection of Cu^2+^-mediated DNA degradation system. DNA protection assays with GroEL1/GroEL1Δ18/*E. coli* GroEL were undertaken. Oxidation of ascorbic acid in presence of cupric ion produces cuprous ion which in presence of hydrogen peroxide releases hydroxyl radicals. These hydroxyl radicals are known to damage DNA. pBSK plasmid DNA (400ng) was incubated with 100 μM CuCl_2_, 1 mM ascorbic acid and 10 mM hydrogen peroxide for 10 min at room temperature in the absence or presence of GroEL1/GroEL1Δ18/ *E.coli* GroEL. Samples were separated on 1% agarose gel and stained with ethidium bromide. Result suggests that GroEL1 protects DNA damage.

## Discussion

Mycobacterial GroEL1 was shown to be lower oligomer according to size exclusion chromatography using molecular weight standards (25). When we characterized the purified GroEL1 using size exclusion chromatography coupled to multi angle light scattering (SEC-MALS), we found that molecular weight was approximately equal to the theoretical molecular weight of a monomer (Figure 2). Next, we removed the 18 residues Histidine-rich C-terminal and again performed SEC-MALS. The flexible C-terminal may affect the oligomerization (Figure 1) however, it remained as a monomer (Figure 2). It is reported in *E. coli* GroEL that the equatorial domain is responsible for oligomerization (26). We purified GroEL1 equatorial domain containing the extremities (1-133 and 409-539 residues) of the polypeptide sequence, connected by the linker (serine, glycine and serine residues). SEC-MALS analysis of the GroEL1 equatorial domain showed molecular weight to be a monomer (Figure S1). SEC based molecular weight determination is based on the hydrodynamic radius of the protein which is estimated relative to the molecular weight standards whereas SEC-MALS does not rely on the standards but light scattering to calculate the absolute molecular weight of the protein. GroEL1 belongs to the family of proteins called chaperonins that needs to be tetradecameric to carry out its protein folding mechanism (36). These lower oligomer may be responsible to carry out some other functions (moonlighting) (37). We also have earlier reported that GroEL1 may act as nucleoid associated protein (35).

Due to the presence of the characteristic Histidine-rich C-terminal in GroEL1 contrary to (GGM)_4_ repeat in its paralog GroEL2 and *E.coli* GroEL (28), we tried to decipher the functional role of these last 18 residues. It is reported that GroEL1 binds nickel and functionally it is known to bind iron for biofilm formation in *Mycobacterium smegmatis* (28). In another study from our lab, it is reported that *M. tuberculosis* GroEL1-KO (knock out) strain is sensitive under low aeration stress and 9 of the 30 copper regulated genes (20) are down regulated (38). By performing ITC studies we found that GroEL1 binds copper (Figure 3A) and has two binding sites (Table 1). It also binds nickel and cobalt but with lower affinity and has one binding site (Table 1). It did not show binding to iron (Figure 3D). Upon deletion of the 18 Histidine rich C-terminal residues (GroEL1-Δ18), the metal binding property was lost (Figure 4A). To identify the specific domain of GroEL1 protein which binds copper, we purified GroEL1 equatorial domain (GroEL1-ED) (Figure 1). The ITC analysis with this domain confirmed copper binding. The weak endothermic binding isotherm was lost suggesting that the equatorial domain is responsible for copper binding and the weak binding site (observed in GroEL1 and GroEL1-Δ18) is not in the equatorial domain (Figure 4B). These results indicate that the Histidine-rich C-terminal of GroEL1 binds copper.

To address the mechanistic effect of copper binding to GroEL1 protein on the overall shape, we performed SAXS studies on GroEL1 in absence and presence of copper in the molar ratio of 1:2, 1:5 and 1:10. The SAXS profile of GroEL1 protein in presence of copper show some differences and it is evident in the structure model that suggests that the structure opens up and flexibility is increased (decrease in Porod Exponent value). The increase in R_g_ value does not hint at oligomerization since the molecular weight estimated through Vc (Volume of correlation) does not add up to a dimer or higher oligomer. It is quite evident that there is increase in size of the protein by opening up (Figure 6, Table 3). This was confirmed by monitoring ANS dye binding to GroEL1 protein in presence of copper which measures extent of hydrophobic residues binding to protein due to the conformation change. Addition of copper (Cu^2+^) ions to GroEL1 protein causes decrease in emission intensity of the dye (Figure S2). Copper chaperones play important role in shuttling metal ions to target proteins. The structure of metallochaperones must be sufficiently flexible both to carry specific ions while cytoplasmic movement and to transfer the ions to or from a partner protein (39). SAXS studies on GroEL1 protein show that GroEL1 protein is flexible and binding of copper (Cu^2+^) ions causes conformational change in GroEL1 as well as increases its flexibility. Our data provide evidence that conformational changes accompany binding of copper to GroEL1.

Once it was confirmed that GroEL1 binds copper, we tried to understand functional implication of GroEL1 and copper interaction in *M. tuberculosis groEL1*-KO (knockout) strain. It is reported that copper is an important micronutrient whose homeostasis is essential, otherwise highly toxic when present at higher concentrations (20). The first insight into the role of copper at the host-pathogen interface was reported by the finding that copper level fluctuated from 25 to 500 µM in the phagosome infected with *M. tuberculosis* (4). Recently it has been reported that utilizing abundance of copper ions in the *M. tuberculosis* infected phagosome using peptide that binds copper ions and mediates generation of reactive oxygen species by exploiting the redox property of copper ions (40). We used mycobacterial strains containing *groEL1*-WT (wild type), *groEL1*-KO (knockout) and *groEL1*-COMP (complemented) gene to study effect of copper on survival at different concentrations. It was observed that *groEL1*-KO is sensitive to 100 µM copper concentration in the culture media (Figure 7). Thus, GroEL1 appears to protect *M. tuberculosis* from the toxic effects of copper (Cu^2+^) ions.

In order to understand the physiological implications of GroEL1 under high copper stress, we undertook biochemical studies. Earlier, it was reported that there was a strong correlation between the copper induced transcriptome and the genes induced by oxidative stress (20). Copper is known to be a redox-active metal and under physiological conditions, cycles between two oxidation states of Cu^1+^ and Cu^2+^ ions. Its redox potential enables catalysis of oxidation reactions using molecular oxygen (12). Hence, it becomes necessary to maintain redox balance against higher copper concentrations. We employed ascorbic acid as a model substrate to study the reduction of cupric (Cu^2+^) to cuprous (Cu^1+^) state. However, in presence of GroEL1, this phenomenon diminished proving that GroEL1 may be involved in redox silencing inside the cell (Figure 8) also we can also say that GroEL1 does not facilitate the reduction of Cu^2+^ to Cu^1+^, which is oxidation state in which copper is exported and hence it may be helpful in transferring Cu^2+^ to important enzymes for its activity or to transcription factors for regulating gene expression. Copper is also known to take part in Fenton-type reaction in presence of hydrogen peroxide to generate hydroxyl radicals (41). Hydrogen peroxide is a by-product of aerobic respiration (42). The hydroxyl radicals are known to react with most of the macromolecules, including nucleic acids (14, 15). Since GroEL1 is known to act as a nucleoid associated protein in *M. tuberculosis* (35), we tried to find out whether GroEL1 helps in protection from Cu^2^-catalyzed DNA damage by hydroxyl radicals. Indeed, in the presence of GroEL1, there was protection of DNA when incubated along with ascorbic acid, cupric chloride (CuCl_2_) and Hydrogen peroxide (H_2_O_2_) (Figure 9).

These results revealed that *M.tuberculosis* GroEL1 has a novel property of maintaining copper homeostasis when copper is present in excess and its deletion resulted in increased sensitivity to copper stress. It may act as a copper chaperone indicated in other bacteria such as in *Kineococcus radiotolerans* which is shown to be copper resistant and possesses GroEL paralog having Histidine-rich C-terminal (43). We propose that the mode of action could be GroEL1 binding Cu^2+^ ions and mediating in redox balance and protection from hydroxyl radical mediated DNA damage (Figure 10). Copper having redox active states i,e Cu^2+^ and Cu^1+^, generates hydroxyl radicals undergoing Haber-Weiss and Fenton-type reactions. The model on the basis of *in vitro* studies with GroEL1 protein show that by binding to Cu^2+^, it reduces the formation of Cu^1+^ ions and downstream generation of hydroxyl radicals. The model depicts the copper (Cu^2+^) ions sequestration mechanism of GroEL1 protein.

**Figure 10:**
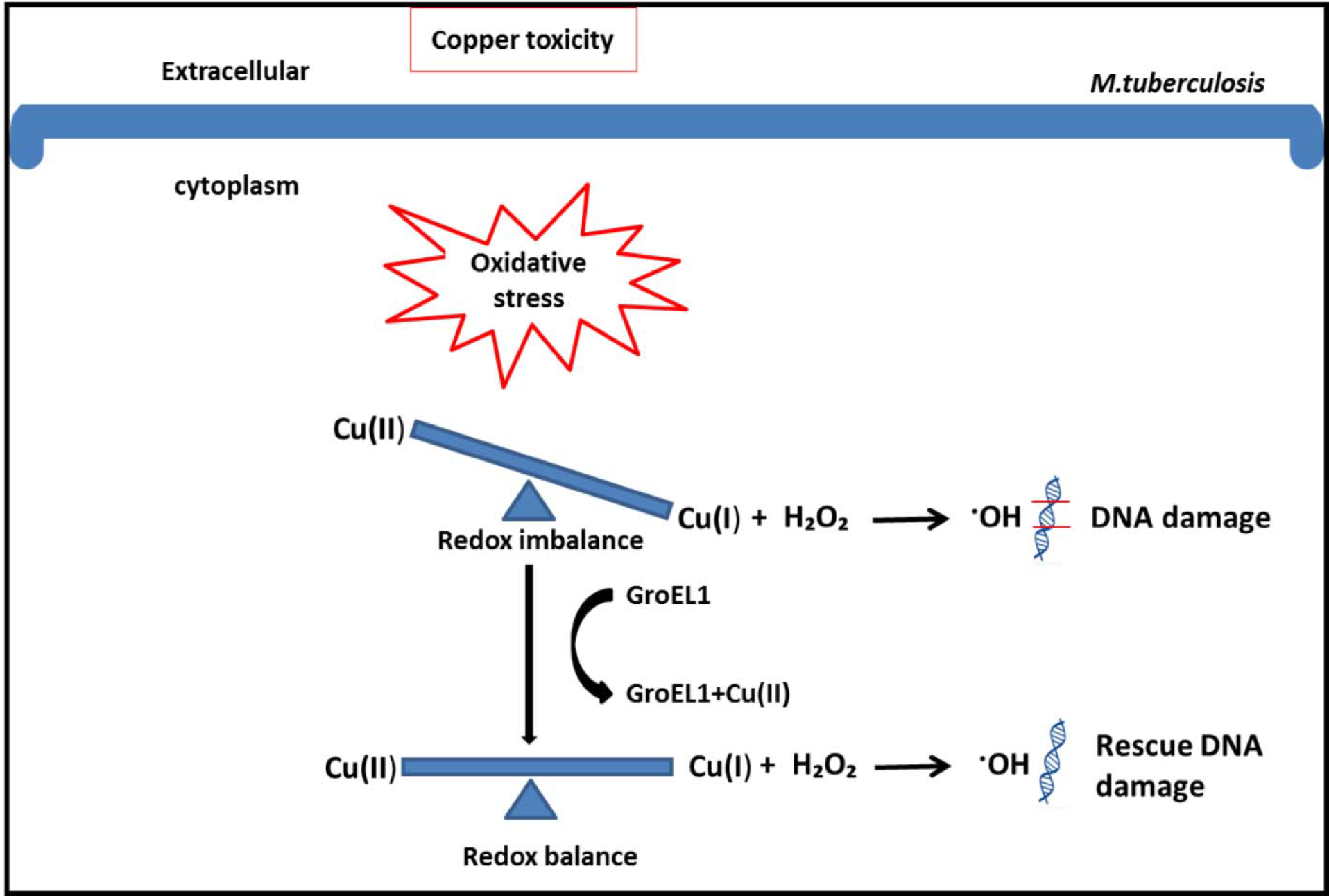
Model for GroEL1 mediated copper homeostasis in *M. tuberculosis*. *M. tuberculosis* under high copper concentration is known to induce oxidative stress (Ward et al., 2008). Copper is a redox-active metal and cycles between two oxidation states, Cu^2+^ and Cu^1+^ ions. Copper is also known to take part in Fenton-type reaction in presence of hydrogen peroxide to generate hydroxyl radicals. We propose on the basis of *in vitro* studies that the *M. tuberculosis* GroEL1-KO (knockout) strain is sensitive to copper toxicity and purified GroEL1 protein binding to Cu^2+^ ions mediates in redox balance and protection from hydroxyl radicals mediated DNA damage.

## Experimental procedures

### Sequence analysis and Homology modeling

Gene sequences of *M. tuberculosis groEL1*, *groEL2* and *E.coli groEL* having characteristic C-terminal were analysed in Jalview (44). The multiple sequence alignment was done using Clustal Omega (45). Homology model of *M. tuberculosis* GroEL1 was built using chain A, crystal structure of Cpn 60.2 from *M. tuberculosis* at 2.8 Å resolution (PDB ID, 3RTK A). The software used was Modeller 9.18v (46). Structure model was validated using Verify 3D (47, 48). Secondary structure prediction was done using PSIPRED (49).

### Plasmids, Protein expression and purification

*M. tuberculosis* full length *groEL1* gene was cloned in pkk223-3 expression vector and designated *groEL1*-full length. The *groEL1* C-terminal 18 residues deletion construct designated *groEL1*-Δ18 and *groEL1* equatorial domain containing construct designated *groEL1*-ED (1-131 and 405-523 residues of the polypeptide sequence, connected by the linker having serine, glycine and serine residues) were previously cloned in our laboratory in the expression vector pET23a. All the constructs were without any affinity tag. The expression vectors were transformed into *E. coli* BL21 (DE3) strain. The culture grown at 37°C at 200 rpm until O.D @ 600 nm reached 0.2 – 0.3. The expression was induced by 1 mM IPTG. The culture was allowed to grow at 18°C at 200 rpm for 12 hours. The pellet obtained after harvesting was resuspended in buffer containing 50 mM Tris-Cl pH 8, 1 mM EDTA pH 8, 150 mM NaCl and Roche protease inhibitor cocktail. The cells were then lysed by sonication (QSONICA Sonicators). The Lysate was centrifuged to obtain supernatant containing protein using Sorvall Centrifuge.

The purification of GroEL1 and GroEL1-ED were optimized using Ni-NTA affinity chromatography taking into consideration the characteristic Histidine rich C-terminal of GroEL1 (Ojha et al., 2005). The supernatant was loaded onto a Bio-Rad disposable plastic column containing Qiagen Ni-NTA resins (5ml) which were washed first with Milli-Q water and then equilibrated with binding buffer (50 mM Tris-Cl pH 8.0, 150 mM NaCl). This was kept on rocker at 4ºC for 1 hour for batch-binding. The beads were then washed with wash buffer (50 mM Tris-Cl pH 8.0, 150 mM NaCl and 10 mM imidazole) with 20 column volume. The protein was eluted with 10 column volumes of elution buffer (50 mM Tris-Cl pH 8.0, 150 mM NaCl and 150 mM imidazole. The elution fractions containing protein were dialyzed against dialysis Buffer (50 mM Tris-Cl pH 8.0, 1mM EDTA and 150 mM NaCl).

In case of GroEL1-Δ18 which has Histidine rich C-terminal deleted, was subjected to ammonium sulphate precipitation. Volume equivalent to 30 % ammonium sulphate solution was slowly added to the supernatant obtained after sonication and centrifugation. It was kept for mixing at constant stirring at 4ºC for one hour to ensure complete precipitation by ammonium sulphate and then centrifuged. The pellet containing the salted out protein was resuspended and dialyzed against 50 mM Tris-Cl pH 8 and 1mM EDTA pH 8. The dialyzed protein was loaded over an anion exchange column, QFF using a Fast protein liquid chromatography (FPLC) system (Bio-Rad laboratories), previously equilibrated with equilibration buffer (50 mM Tris-Cl pH 8 and 1 mM EDTA pH 8) at 4°C. It was subjected to an isocratic flow of 10 column volume with wash buffer (50 mM Tris-Cl pH 8, 1 mM EDTA pH 8 and 150 mM NaCl), followed by 10 column volume of elution buffer (50 mM Tris-Cl pH 8, 1 mM EDTA pH 8 and 300 mM NaCl). The elution fractions containing protein were dialyzed against Dialysis Buffer (50 mM Tris-Cl pH 8, 1 mM EDTA pH 8 and 150 mM NaCl). The dialyzed protein were further passed through size exclusion chromatography using Superdex 200, 16/600 column (GE Healthcare Life Sciences), pre-equilibrated with 50 mM Tris-Cl pH 8, 1 mM EDTA pH 8 and 150 mM NaCl. The peak fractions were pooled and concentrated and finally dialyzed against 10 mM Tris-Cl pH 8. Protein estimation was done using Bradford reagent (Sigma Inc.) and stored in −80°C after flash freezing in liquid nitrogen.

### Size exclusion chromatography coupled multi angle light scattering (SEC-MALS)

Purified GroEL1, GroEL1-Δ18 and GroEL1-ED were subjected to SEC-MALS analysis. It was performed in buffer containing 10 mM Tris-Cl pH 8 and 150 mM NaCl. The protein was dissolved in the same buffer to make different concentrations. SEC-MALS measurements were done using Superdex 200, 10/300 column (GE Healthcare Life Sciences) connected to Agilent HPLC system coupled to a Wyatt Technologies Dawn Heleos II multi-angle light scattering detector and Optilab T-Rex refractometer. BSA (Pierce Albumin Standard, Thermo Fisher Scientific) solution (2 mg/ml) was used to prepare a method after adjusting delay volume, baseline and band broadening parameters. Protein samples were loaded on to the column, varying concentrations from 1.5 mg/ml to as high as 10 mg/ml. The data were analyzed using the ASTRA software (Wyatt Tech). The refractive index increment, d*n*/d*c* of 0.185 was used for the calculation of protein molecular weight.

### Isothermal titration calorimetry (ITC)

ITC experiments were performed using the MicroCal iTC_200,_ (GE). The GroEL1, GroEL1-Δ18 and GroEL1-ED proteins were dialyzed extensively in 10 mM Tris-Cl pH 8. Metal salts of 1M stock concentration were prepared in HPLC grade water and working concentration of metal salts were prepared in 10 mM Tris-Cl pH 8. Approximately 25 µM of GroEL1, GroEL1-Δ18 and GroEL1-ED were used in the experiment. Protein was kept in the cell. 1 mM of chloride salts of Cu^2+^, Fe^2+^, Fe^3+^, Zn^2+^, Mg^2+^, Ni^2+^, Co^2+^ and Ca^2+^ were used in the experiment and kept in the syringe. The experiments were performed at 25 °C with constant stirring of syringe set at 1000 rpm. Titration with metal salts was initiated by injecting 0.4 μl followed by 18 successive 2 μl injections. The time between successive injections was kept at 150 s. Injections of metal salts into reaction buffer in the cell without proteins were performed as controls. The integrated heats of reaction from each injection normalized to the moles of ligand per injection were fit to different binding model using ORIGIN software (Origin Lab, Northampton, MA). The integrated peak of the first injection was excluded from Data fitting analysis.

### Small angle x-ray scattering (SAXS)

#### Data collection

Purified protein samples were subjected to small angle X-ray scattering (SAXS) at ESRF (Grenoble, France) on BioSAXS beamline BM29 (50). The scattering data for each sample and its matching buffer was collected on Pilatus 1M detector (Dectris) in ten frames, 0.5 second exposure per frame at 277 K temperature in ‘flow mode’ using a standard set up (51). The X-ray beam of wavelength λ = 0.991 Å, sample to detector distance of 2.87 meters, scattering vector range (Q) of 0.035 to 0.49 nm^−1^ are the data collection parameters employed at beamline (Table 2). The protein sample was re-suspended in 10 mM Tris-Cl pH 8 for making different concentration range (5mg/ml and 2.5mg/ml). The same buffer was used for buffer subtraction. GroEL1 protein and its complex with copper (Cu^2+^) in the 1:2, 1:5 and 1:10 molar ratio were exposed to X-ray beam for scattering data collection. The data collected for the buffer and the samples were processed and subtracted automatically using EDNA based processing pipeline at ESRF, BM29 (52). Further analysis was done using the PRIMUS software (53) available in the ATSAS 2.8 package suite (54). Scattering intensity I(Q) as a function of scattering vector Q (Q = 4πsinθ/λ, θ is scattering angle) plot was obtained. All the processing conditions were same for all the datasets collected. The I(Q) profile of the buffer was subtracted from the I(Q) profile of the sample solution to obtain the I(Q) profile of the protein molecules in solution.

**Table 2:** SAXS data collection parameters employed during data collection

| **Data-collection parameters** |  |
| --- | --- |
| Beamline | BM29 |
| Detector | Pilatus 1M |
| Wavelength ($\lambda$), Å | 0.991 |
| Detector-to-sample distance (m) | 2.867 |
| Q-range ($\text{nm}^{-1}$) | 0.035-4.94 |
| No. of frames collected | 10 |
| Exposure time<br>Per frame (second) | 0.5 |
| Measurement<br>Temperature (K) | 277 |

**Table 3:** Structural parameters deduced from SAXS data. Structural parameters deduced from SAXS data analysis of GroEL1 protein and Cu^2+^ ions (CuCl_2_) complex in the molar ratio 1:2, 1:5 and 1:10. Concentration of GroEL1 protein was estimated using Bradford assay.

|  |  | **Apo GroEL1**<br>(Concentration-range) |  | **GroEL1-Copper**<br>(1:2 molar ratio) | **GroEL1-Copper**<br>(1:5 molar ratio) | **GroEL1-Copper**<br>(1:10 molar ratio) |
| --- | --- | --- | --- | --- | --- | --- |
| Concentration range (mg/ml) |  | 5 | 2.5 | 5 | 5 | 5 |
| Concentration range ($\mu\text{M}$) | | 90 | 45 | 90:180 | 90:450 | 90:900 |
| **Structural parameters** |  |  |  |  |  |  |
| Guinier Analysis | $R_g$ (nm) | 3.32 | 3.21 | 3.28 | 3.74 | 5.07 |
| | $R_c$ (nm) | 1.7 | 1.67 | 1.7 | 1.7 | 1.7 |
|  | L (nm) | 9.88 | 9.5 | 9.72 | 11.54 | 16.55 |
| Indirect Fourier Transformation P(r) | $D_{\text{max}}$(nm) | 12.49 | 12.36 | 12.63 | 13.59 | 19.75 |
| | $R_g$ (nm) | 3.81 | 3.88 | 3.89 | 4.02 | 5.26 |
| Porod Exponent Estimate |  | 3.5 | --- | 3.4 | 3.3 | 3.2 |
| **Molecular-mass determination (kDa)** |  |  |  |  |  |  |
| Molecular mass $M_r$ [from $V_c$] | | 53.4 | 58.5 | 56.3 | 62.8 | 80.1 |
| Monomeric $M_r$ [from sequence] | | 55.8 | 55.8 | 55.8 | 55.8 | 55.8 |
| **Ab-initio modelling (GASBOR)** |  |  |  |  |  |  |
| Chi-square ($\chi^2$) | | 57.08 | --- | 9.05 | 4.16 | 6.96 |

#### Structural parameters

The double log plot (Log_10_ I(Q) versus Log_10_ Q) was generated to analyze the monodispersity and lack of aggregation in each dataset. Kratky plots (Q versus I(Q).Q) for each dataset estimated the globular or extended nature of the protein and its complex with copper (Cu^2+^). The Guinier analysis (Q2 versus Ln I(Q) was done using the low Q-region to estimate radius of gyration (R_g_) and radius of cross-section (R_c_), assuming globular and rod-like scattering shape of the predominant scattering molecules in solution. Using the equation L={12(R_g_^2^-R_c_^2^)}^1/2^, persistence length of the respective scattering entities were estimated. Further, indirect Fourier transformation of each dataset was done over a measured Q-range to obtain a pairwise distance distribution function P(r) of interatomic vectors (55). The probability of finding the pairwise interatomic vectors at 0 Å length and maximum linear dimension (D_max_) of the protein molecules in solution was considered to be zero. A plot of Porod Exponent was generated to extrapolate the conformational changes occurring in presence of ligand copper (Cu^2+^) (30). Molecular weight was estimated based on the volume of correlation (V_c_) using DATVC program in ATSAS 2.8 suite.

#### Model building

*Ab initio* chain-ensemble models of GroEL1 protein and its complex with copper (Cu^2+^) were generated using the GASBOR program. Using 540 dummy residues (considering GroEL1 as monomer based on SEC-MALS data), with no shape and symmetry bias, low resolution structures were reconstructed within the shape constraints derived during distance distribution function P(r) analysis. Model building of each dataset was repeated five times. Theoretical I(Q) profile of these five models were compared with I(Q) profile of the raw data using CRYSOL program and the model with the lowest Chi-square (χ^2^) value was selected as final low resolution shape of the GroEL1 protein and its complex with copper (Cu^2+^). The resultant model of GroEL1 and its complex with cupric ion was aligned in automated manner with the Homology model of GroEL1 protein generated using Modeller software (46) using SUPCOMB program. All the programs used are part of the ATSAS 2.8 suite. PyMOL, GraphPad Prism and ORIGIN were used for figure generation and graphical analysis.

### ANS (8-anilino-1-naphthalenesulfonate) binding assay

ANS (8-anilino-1-naphthalenesulfonate) binding experiments were done using Agilent Cary Eclipse Fluorescence Spectrophotometer and cuvette of 1cm path-length. Fluorescence spectra were collected at excitation wavelength of 370nm and emission wavelength at 400-600 nm range. The ANS fluorescent intensity of of GroEL1 (apo) protein and in presence of copper (Cu^2+^) in the molar ratio of 1:5, 1:10 and 1:20 were recorded at room temperature. The concentration of GroEL1 and ANS were 20 μM and 100 μM respectively. GroEL1 and copper (Cu^2+^) mixture were first incubated for 2 hours and then further incubated for 1 hour after adding ANS. It was done in tubes covered in aluminium foil and at room temperature. Control experiments include recording fluorescence spectra of ANS alone as well as in presence of copper without adding protein.

### Mycobacterial viability assay

The *M. tuberculosis* H37Rv strains containing *groEL1* wild type gene (*groEL1-*WT), *groEL1* knockout gene (*groEL1*-*KO*) and *groEL1* complemented gene (g*roEL1-*Comp) were obtained from the lab of Dr Anil Ojha (28). These strains were cultured in 7H9 supplemented with OADC and 0.05% tyloxapol to A_595_ ~0.8-1. The strains were then sub-cultured in the same medium and grown to an A_595_~0.8. These secondary cultures were used for the experimental set up. Briefly, the cultures were washed in 7H9 self-made medium (*i.e.* without CuSO_4_ + 0.2% glycerol + 0.05% tyloxapol) and diluted to A_595_~ 0.15-0.2 in 7H9 self-made medium. The diluted cultures were dispensed in 10 ml aliquots each in 50 ml falcon tube. The following conditions were set up in triplicates: Untreated (UT, 0 concentration), 5, 50 and 100 μM CuSO_4._ The effect of addition of CuSO_4_ on the viability of M. tuberculosis strains was monitored over a period of 16 days. At each time point, culture aliquots were taken to monitor A_595_ and ten-fold serial dilutions were plated on 7H11 Agar plates supplemented with 0.5% glycerol and OADC by track-dilution method. Colonies were finalized by 6 weeks.

### Cu^2+^-mediated ascorbic acid oxidation assay

For the oxidation experiments, the ascorbic acid concentration of 300 µM was taken. The absorbance of ascorbic acid was monitored at 260 nm at room temperature using UV/VIS spectrophotometer (BECKMAN COULTER). Quartz cuvette (BECKMAN) of 1 cm path length was used. The oxidation of ascorbic acid is a function of reduction of Cu^2+^ to Cu^1+^ in presence of oxygen. The oxidation of ascorbic acid in 10 mM Tris-Cl pH 8 buffer in presence or absence 5 µM CuCl_2_ was measured at 260 nm during the span of 30 minutes. To study the effect of GroEL1 in binding to CuCl_2_ and inhibiting the oxidation of ascorbic acid, 5 µM of GroEL1 protein was added in the reaction mixture. GroEL1-Δ18 was used as control experiment. The blank reading was taken using the reaction mixture without ascorbic acid. Experiments were done in three technical replicates.

### Cu^2+^-mediated DNA damage assay

Cu^2+^ mediated formation of hydroxyl radicals in presence of ascorbic acid and hydrogen peroxide undergoing Fenton-like reaction can damage DNA. pBSK plasmid was used in this study. 400 ng of plasmid was incubated with 1 mM ascorbic acid, 100 µM CuCl_2_ and 10 mM hydrogen peroxide in 10 mM Tris-Cl pH 8 buffer for 10 min at room temperature. Reaction was stopped by adding DNA loading dye. Status of DNA damage was analysed on 1% agarose gel, stained with ethidium bromide. Role of GroEL1 was checked by adding it in the reaction mixture. GroEL1-Δ18 and *E.coli* GroEL were used as control. Experiments were performed in three replicates.

## Acknowledgements

The funding was provided by Department of Biotechnology, Ministry of Science and Technology grants BT/PR3260/BRB/10/967/2011 and BT/PR15450/COE/34/46/2016. We acknowledge ESRF-DBT for funding ESRF-Grenoble visit for SAXS data collection, Beamline staff at the ESRF (XRD1) for their assistance during data collection. MYA also acknowledge Department of Biotechnology for fellowship and Kriti Chopra (NCCS-Pune) for helping in homology modeling of GroEL1 protein.

## Conflicts of interest

The authors declare that they have no conflicts of interest with the contents of this article.

## Author contributions

MYA, SCM designed the study; MYA performed all the experiments; SDB performed the M. tuberculosis H37Rv Copper toxicity experiments and SDB, JST analysed the data; HO helped in ITC experiments, Dr Ashish helped in SAXS data collection and analysis; MYA, SCM wrote the manuscript, Dr Ashish, SDM and JST wrote specific part of manuscript. SCM obtained the funding and supervised the project. All authors have read the manuscript

## Abbreviations

M. tuberculosis: Mycobacterium tuberculosis
E. coli: Escherichia coli
ITC: isothermal titration calorimetry
SEC-MALS: size exclusion chromatography coupled multi angle light scattering
ANS: 8-anilino-1-naphthalenesulfonate
ASA: ascorbic acid
pBSK: pBluescript SK (-) plasmid

