## SUPPLEMENTARY MATERIAL for "A novel function of *M. tuberculosis* chaperonin paralog GroEL1 in copper homeostasis"

**Running title:** *M. tuberculosis* GroEL1 involved in copper homeostasis

[χ]**Present address:** Council of Scientific and Industrial Research (CSIR), Anusandhan Bhawan, 2 Rafi Marg, New Delhi-110001, INDIA.

[Ψ]**To whom correspondence should be addressed**:
Shekhar C. Mande
National Centre for Cell Science, NCCS Complex,
SP Pune University Campus, Ganeshkhind
Pune- 411 007, INDIA



**keywords:** *Mycobacterium tuberculosis,* GroEL1, Gene duplication, Copper homeostasis, redox balance

**LIST OF MATERIALS INCLUDED:**

**Figure S1**
**Figure S2**
**Figure S3**
**Figure S4**

Supplementary figure 1

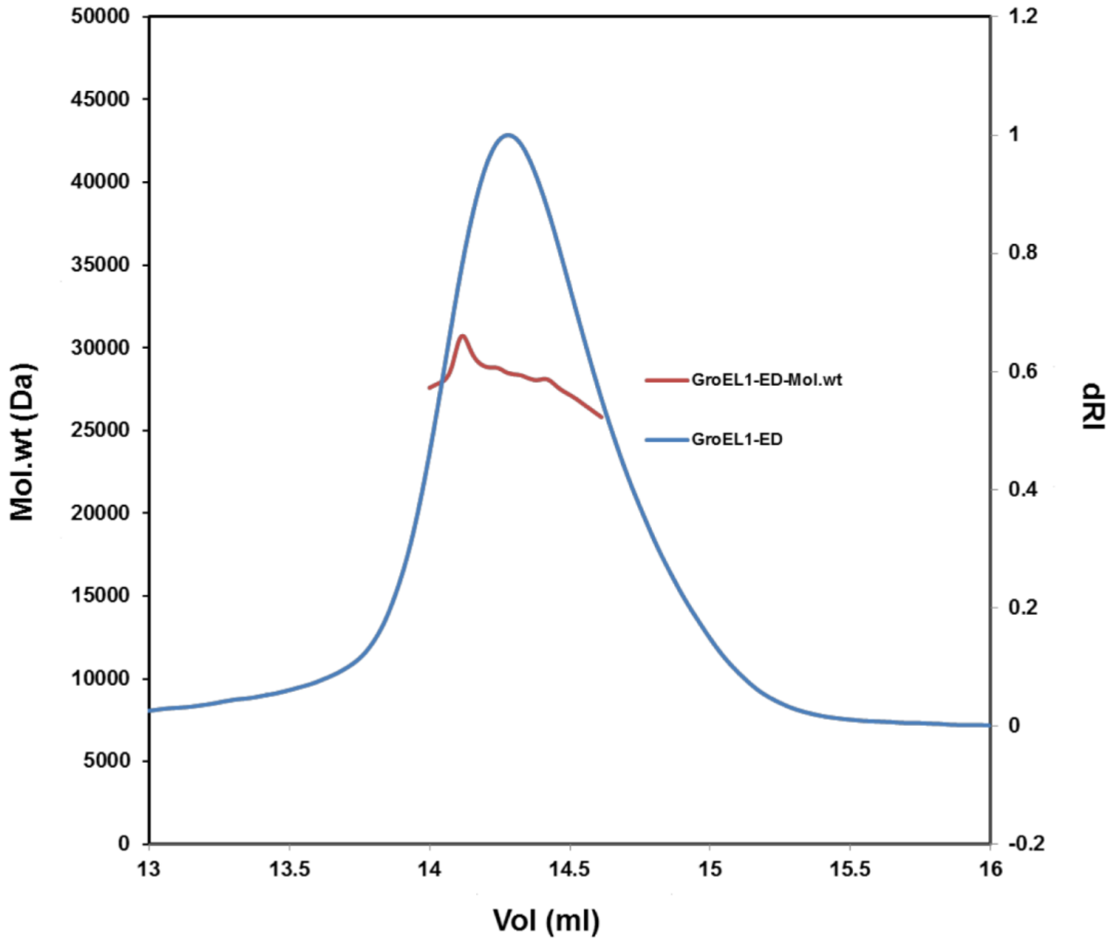


**Figure S1: GroEL1-ED (equatorial domain) exists as monomer as shown by SEC-MALS.**

SEC-MALS elution profiles of GroEL1-ED in 10 mM Tris-Cl pH 8.0 and 150 mM NaCl at 7 mg/ml concentration. The molecular mass distribution is shown as thick continuous line. The observed molecular weight using Astra software (Wyatt-Tech) is approximately equal to the theoretical molecular weight of a monomer i.e. 27 kDa.

Supplementary figure 2

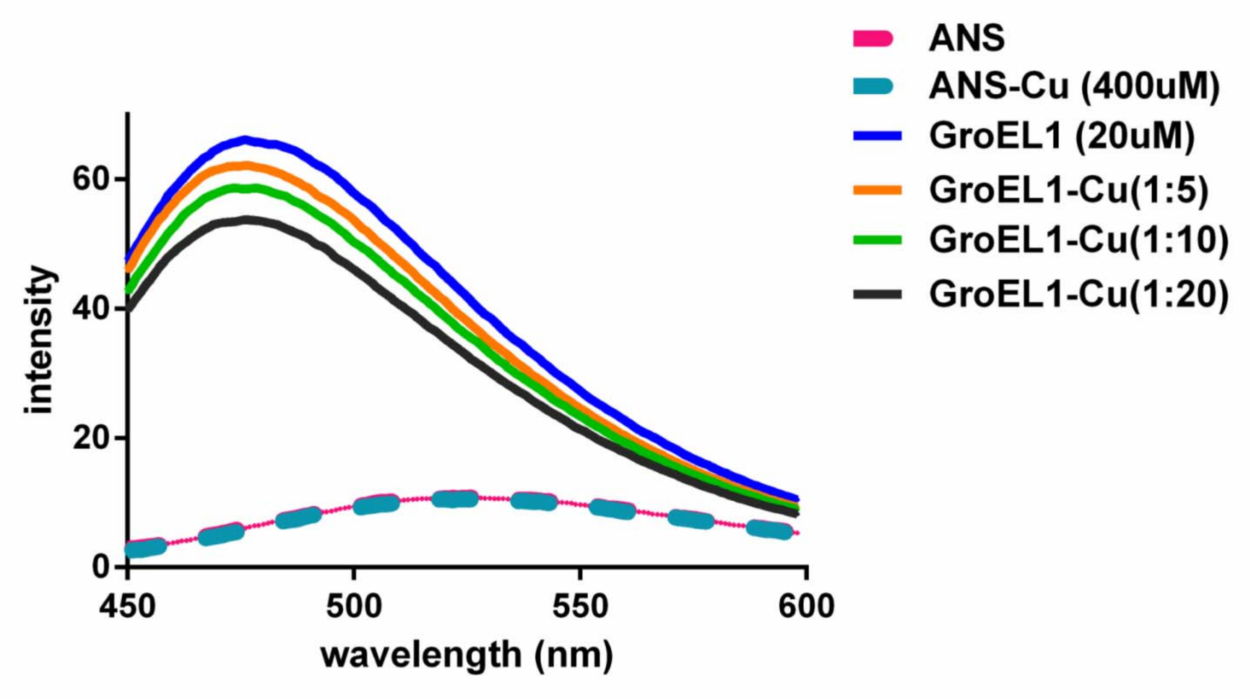


**Figure S2: Monitoring structural change of GroEL1 with copper ($Cu^{2+}$) through ANS binding**

Emission spectra of ANS bound to GroEL1 and its complex with $Cu^{2+}$ ions ($CuCl_2$). Observation of decrease in fluorescence intensity of ANS upon increasing concentration $Cu^{2+}$ ions ($CuCl_2$) in the molar ratio of 1:2, 1:5 and 1:10. Each spectra is a mean of four replicates

Supplementary figure 3

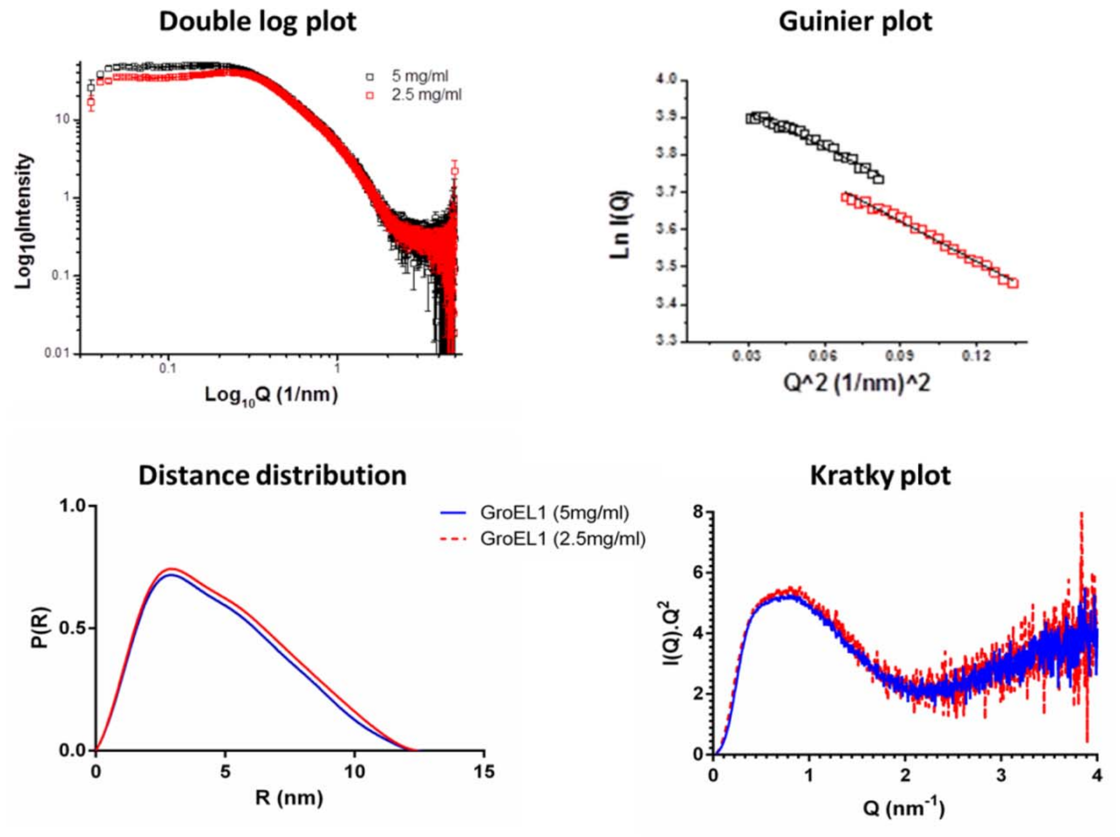


**Figure S3: SAXS data analyses and plots of GroEL1**

SAXS data analysis of GroEL1 protein at concentration of 5 and 2.5 mg/ml. (A) Double-Log plot of SAXS data of GroEL1. (B) Guinier plot of GroEL1 (data curve is modified for ease in visualisation). (C) Pairwise distance distribution plot of GroEL1. (D) Kratky plot of GroEL1.

Supplementary figure 4

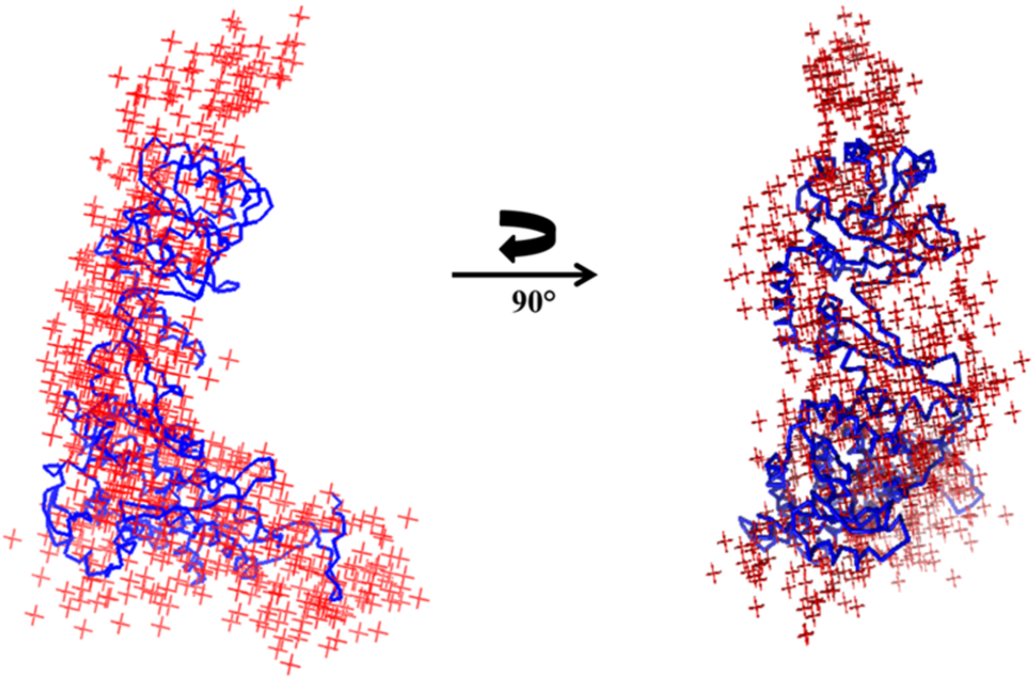


**Figure S4: SAXS based *ab initio* model of GroEL1**

Predominant solution shape of SAXS based model of GroEL1 (5 mg/ml). Figure shows *ab initio* model (GASBOR) of GroEL1 protein in two orientations. The model is superimposed on homology model of GroEL1 protein.
